# Characterizing transition cells in developmental processes from scRNA-seq data

**DOI:** 10.1101/2022.05.18.492572

**Authors:** Yuanxin Wang, Vakul Mohanty, Jinzhuang Dou, Shaoheng Liang, Qingnan Liang, Yukun Tan, Jin Li, Ziyi Li, Rui Chen, Ken Chen

**Affiliations:** Department of Bioinformatics and Computational Biology, The University of Texas MD Anderson Cancer Center; Department of Biostatistics, The University of Texas MD Anderson Cancer Center; Molecular and Human Genetics, Baylor College of Medicine

## Abstract

Multi-cellular organism development involves orchestrated gene regulations of different cell types and cell states. Single-cell RNA-Seq, enable simultaneous observation of cells in various states, making it possible to study the underlying molecular mechanisms. However, most of the analytical methods do not make full use of the dynamics captured. Here, we model single-cell RNA-seq data obtained from a developmental process as a function of gene regulatory network using stochastic differential equations (SDEs). Based on dynamical systems theory, we showed that pair-wise gene expression correlation coefficients can accurately infer cell state transitions and validated it using mouse muscle cell regeneration scRNA-seq data. We then applied our analytical framework to the PDAC (Pancreatic ductal adenocarcinoma) mouse model scRNA-seq data. Through transition cells found in the pancreatic preinvasive lesions scRNA-seq data, we can better explain the heterogeneity and predict distinct cell fate even at early tumorigenesis stage. This suggests that the biomarkers identified by transition cells can be potentially used for diagnosis, prognosis and therapeutics of diseases.

## Introduction

During the development of multi-cell organisms, different cells make their own decisions to different cell types and cell states. Understanding the underlying molecular mechanisms can deliver deeper insights on physiology, morphology and etiology of diseases. Gene expression, a readout of the developmental processes, opens a window to observe these fundamental molecular processes and construct models such as gene regulatory networks (GRNs) to understand differentiations and fate decisions (Cardoso-Moreira et al., 2020). By studying the differentially expressed genes (DEGs) at different stages of the developmental processes, we can identify candidate biomarkers of the process, thus determining the diagnostic signatures and therapeutic targets for diseases (Rodriguez-Esteban & Jiang, 2017).

Traditional ways comparing gene expressions using wet lab experiments leverage molecular biology tools such as real-time quantitative PCR (qPCR). However, qPCR requires specific primers for genes of interest, which limits the discovery power to candidate genes. With the emergence of high-throughput RNA sequencing technique, we can unbiasedly investigate the expression profile of thousands of genes at the same time. Yet, bulk RNA sequencing is only able to detect average expression levels of these genes from cells in different states. The averaging smooth out heterogeneity among these cells, making it hard to characterize the underlying dynamics, especially for state transitions. State transitions, required by differentiation, dedifferentiation and transdifferentiation, play crucial roles in many developmental processes, such as hematopoiesis and tissue regeneration, and can cause diseases if becoming uncontrolled (Brackston et al., 2018; Mulas et al., 2021). Due to a lack of analytical tools, it remains challenging to understand the full pictures of these transitions.

Recently, single-cell RNA-seq has been widely used to study gene expressions of heterogeneous samples. Its ability to measure cell-to-cell variations can reveal complex and rare cell populations, making it possible to study dynamical transition processes (Hwang et al., 2018; Wang et al., 2019). Currently, however, computational tools available for finding cellular states and state transitions are limited. The commonly used methods cluster cells in lower dimensions produced by approaches such as PCA, t-SNE and UMAP, and annotate each cluster based on well-established markers (Hwang et al., 2018). Yet, few reliable markers exist for transition cells comparing to well-defined stable cells. Although trajectory-based methods such as monocle and Slingshot order cells by pseudotime, assuming that state transitions generate continuous expression profiles, they still cannot distinguish transition cells from cells in the stable states, and extract clearsignals to characterize the transition processes (Zhou et al., 2021). While Cellrank (Lange et al., 2022) and Mutrans (Zhou et al., 2021), defines macrostates and attractor basins to separate stable cells and transition cells, rely on the cell-cell similarity without explaining the underlying gene regulatory mechanisms, through, for example, constructing gene regulatory networks (GRNs) (Hwang et al., 2018). Still lacking are systematic ways to discriminate and characterize transition cells from stable cells.

In systems biology, differential equations are popular tools to describe the dynamical processes in living cells. Differential equations typically model evolving gene expressions as rate functions of gene regulatory relations. Model parameters can be interpreted as strength of regulations. After estimating parameters through wet lab experiments or data fitting, we can potentially find numerical solutions and stable/unstable states accordingly (Ioannis Stefanou & Jean Sulem, 2021). However, the approaches have proven to be computationally complex and expensive and can only be applied to model systems involving hundreds of genes. If we want to include more genes to describe the entire developmental processes, the number of variables and parameters become very large, making it challenging for finding solutions and further analysis (Daun et al., 2008; Kreutz, 2020). Instead of trying to decipher the whole regulatory process and underlying regulators, here, we propose that Pearson’s correlation coefficients between gene pairs can be used as the metrics to identify transition cells and understand molecular mechanisms during developmental processes.

## Results

### Modeling gene expressions using SDEs

To model gene expressions as a function of GRNs, we used stochastic differential equations (SDEs) depicting the developmental processes; where **X** denotes gene expression levels, *f*(***X***) a function of regulatory relations, and ***σ****W*_*t*_ the scaled Wiener process:

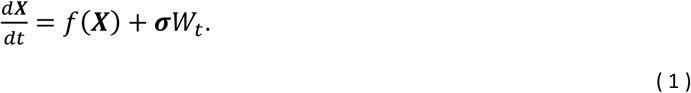

To simplify the model, we can linearize *f*(***X***) using Taylor’s expansion:

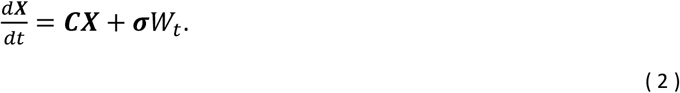

By solving equation (2), the covariance matrix of ***X***_***t***_ can be written as:

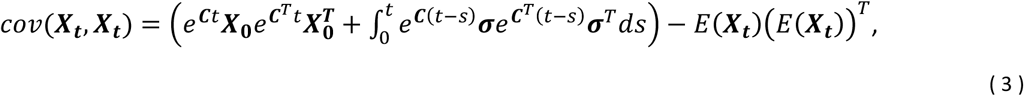

where ***X***_**0**_ is the gene expression levels at the initial time point, while ***X***_***t***_ is the gene expression levels at time *t*. We assume that most of cells captured by scRNA-seq are approximately equilibrium according to Boltzmann distribution. Taking the derivative of the covariance matrix, we arrive at equation (4), the continuous-time Lyapunov equation:

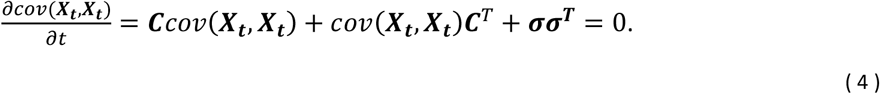

According to Simon et al. (Freedman et al., 2022), one of the conclusions that can be drawn from equation (4) is when a cell is in a transition state, its gene pair-wise Pearson’s correlation coefficients are more likely to be close to ±1. Briefly, the **C** matrix can be diagonalize into ***P*Λ*P***^**-1**^, and equation (4) can be written as 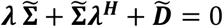, where 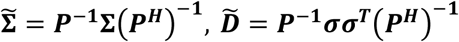. After plugging in the eigenvalues and eigenvectors of matrix **C**, the covariance between gene *i* and *j* can be further simplified as in equation (5):

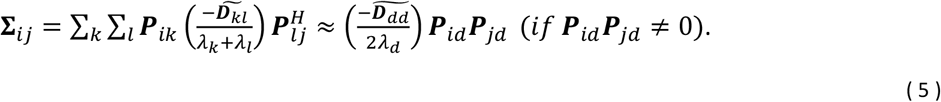

According to bifurcation theory (Ioannis Stefanou & Jean Sulem, 2021), all the eigenvalues of **C** should be negative at stable states, while the maximum eigenvalue (*λ*_*d*_) approaches 0, if the cell is transiting from a stable state to an unstable state:

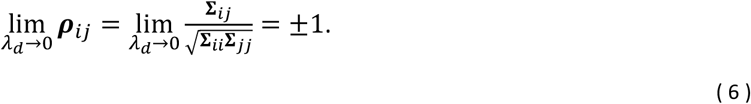

Thus, by using simple metrics, Pearson’s correlation coefficients, we can relate gene expressions with cellular behaviors and identify transition cells during developmental processes.

### Validation using mouse muscle cell regeneration data

To validate our method, we applied our analysis framework to a mouse muscle cell regeneration scRNA-seq data (McKellar et al., 2021), which contain stable cells and transition cells annotated by canonical genes. Because of the small population size of transient cell states, McKellar et al. integrated 111 single-cell RNA-seq datasets to study gene expression dynamics in muscle injury response. Due to the large number of cells in the integrated dataset, we selected a subset of the datasets (**Fig. 1a**) to compare the Pearson’s correlation coefficients between transition cells and stable cells. We calculated gene pair-wise Pearson’s correlations and corresponding transition index for each cell as describe in **Methods**. We found that the transition indices for cells in the transition state are significantly higher than cells in stable states (**Fig. 1b-c**) (Wilcoxon test; p-value < 0.01).

**Fig. 1.**
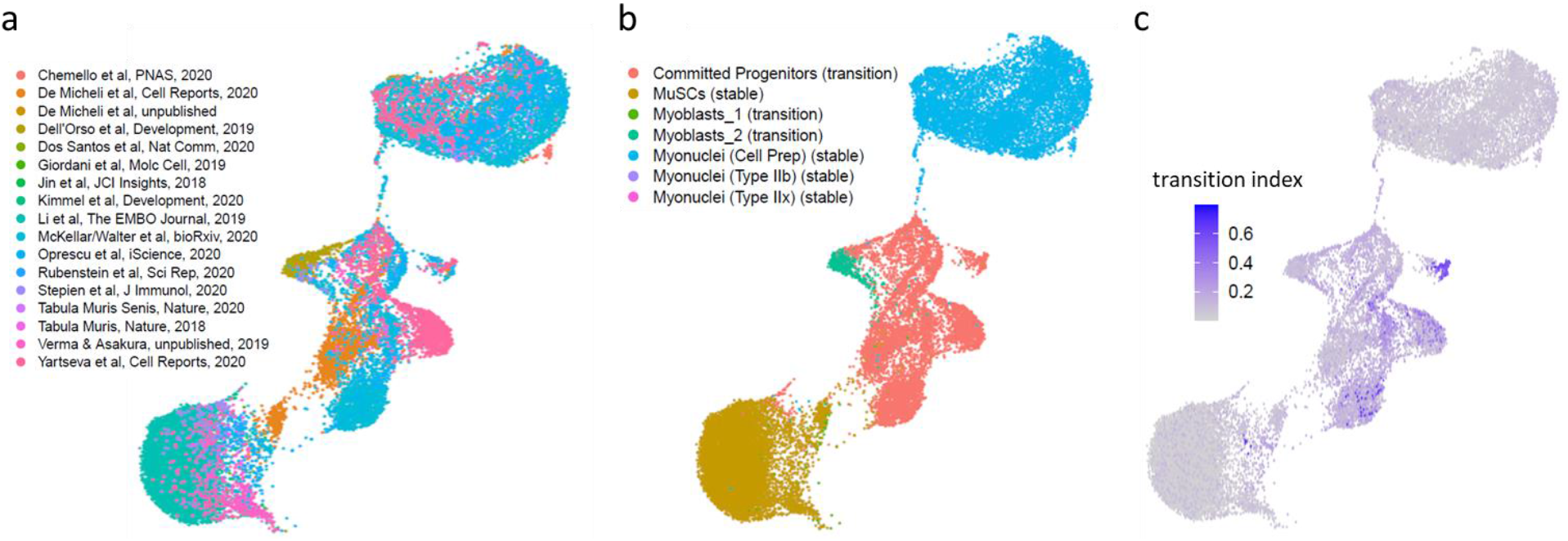
Transition index defined according to the distribution of Pearson’s correlation coefficients can accurately identify transition cells. **a**, A subset of integrated data was selected to validate our method’s capability in identifying transition cells. **b**, UMAP colored by cell types annotated using canonical markers. **c**, UMAP colored by transition index.

### Transition cells in pancreatic preinvasive lesions

We next used our method to investigate the biological processes in pancreatic preinvasive lesions. Pancreatic ductal adenocarcinoma (PDAC) is an aggressive cancer with poor prognoses but has the potential to be cured if being diagnosed at very early stage (*Pancreatic Cancer Prognosis* | *Johns Hopkins Medicine*, n.d.). Schlesinger et al. (Schlesinger et al., 2020) used mouse model to perform a time course scRNA-Seq during the progression from preinvasive lesions to tumor formation, making it possible to explore the early molecular processes giving rise to PDAC. Briefly, they used *Ptf1a-Cre*^*ER*^; *Rosa26*^*LSL-tdTomato*^ mice as control and *Kras*+/*LSL-G12D; Ptf1a-Cre*^*ER*^; *Rosa26*^*LSL-tdTomato*^ mice in the experimental group and injected Tamoxifen to induce preinvasive lesions. Pancreas was collected at 6 time points (17 days, 6 weeks, 3 months, 5 months, 9 months, 15 months post-tamoxifen injection (PTI)) after the injection for sequencing. Metaplastic cells were observed beginning at 3M post-injection (**Fig. 2a**). To study the metaplastic cells during the disease progression, we calculated the eCDF (empirical Cumulative Distribution Function) of the Pearson’s correlation coefficients at different time points (**Fig. 2b**). The transition index is significantly different for cells at different time points (Wilcoxon test; p-value < 0.01). We observed groups of cells in the early stage whose transition indices are higher than cells at the same stage (**Fig. 2c-d**), which suggest that these were the transition cells during the tumorigenesis. To further characterize transition cells and better understand their roles in the early events of preinvasive lesions, we found DEGs of transition cells comparing to other cells at 3M PTI and did a gene set enrichment analysis. Interestingly, transition cells at 3M PTI are heterogeneous and reflect different potential paths of metaplastic cells (**Fig. 2e**). These paths include stomach-like metaplasia and becoming tumor cells, which coincides with the observations at later time points (Ma et al., 2022).

**Fig. 2.**
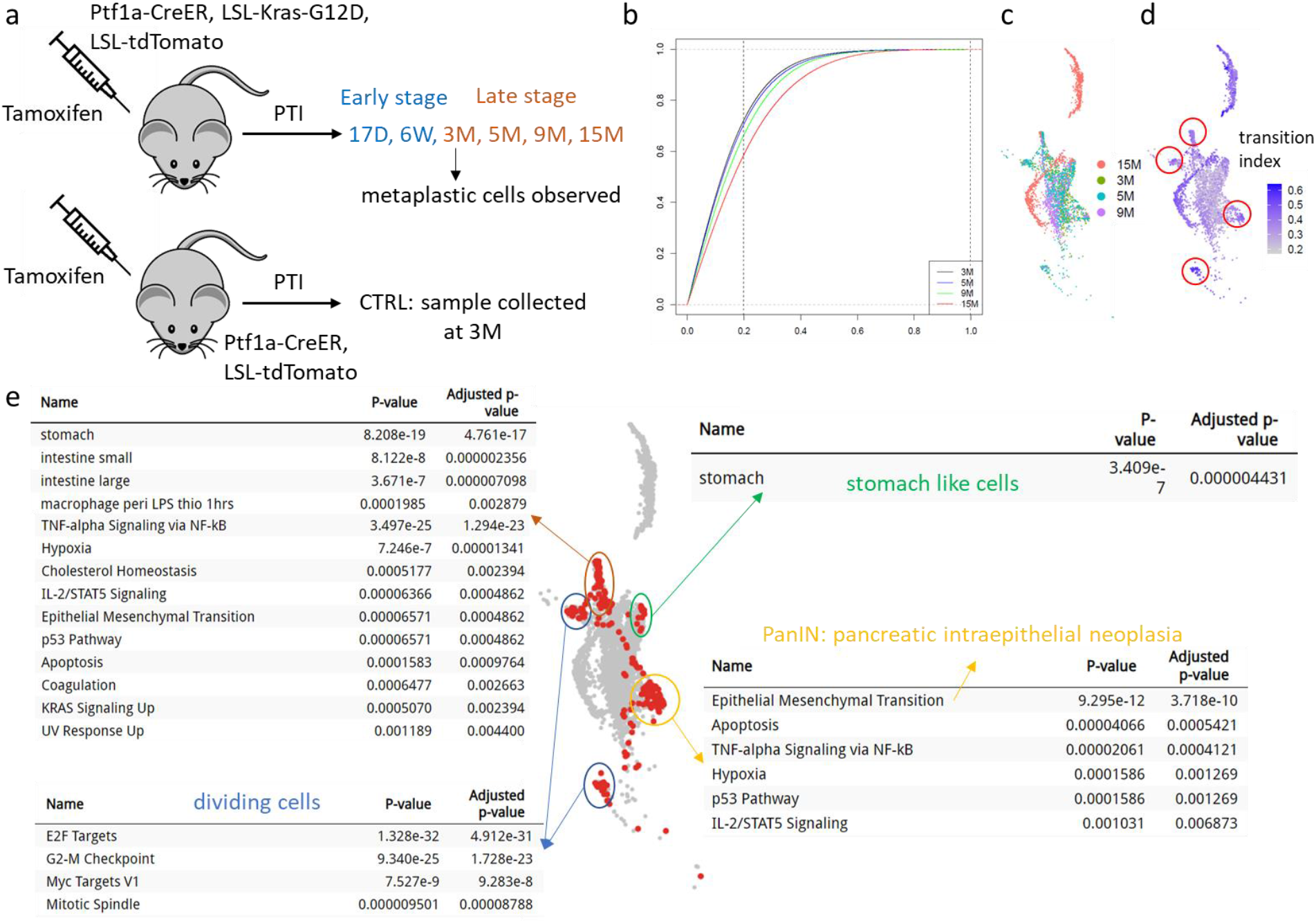
Transition cells reveal the heterogeneity of metaplastic cells even at very early stage. **a**, Study design of the public data GSE141017. **b**, ECDF of gene pair-wise Pearson’s correlations of metaplastic cells at different time points. **c**, UMAP colored by sample collected time points. **d**, UMAP colored by transition index. Red circles: high transition index at early time points **e**, The enrichment analysis of DEGs found using transition cells comparing with baselines at 3M PTI indicates distinct subpopulations of metaplastic cells at later time points.

We then explored the process before the accumulation of pancreatic intraepithelial neoplasia (PanIN) that can be observed through Hematoxylin and Eosin (H&E) staining. The transition index of control groups has no significant difference with that of 17D PTI (Wilcoxon test; p-value=0.34) but not 6W PTI (Wilcoxon test; p-value < 0.01). If we plot the eCDF of gene pair-wise Pearson’s correlation coefficients for each cell, we can see there are two clusters of cells at 6W PTI, and the transition cells accumulate starting from 17D PTI (**Fig. 3b-d**). We found 650 DEGs in total when comparing transition cells at 6W PTI to the control group, where 491 of them are unique for transition cells. Among these 491 genes, many of them are in signaling pathways reported to be deregulated during carcinogenesis (Reyes-Castellanos et al., 2020). And we also found some genes such as *CD47*, a ‘don’t eat me signal’, and *Sox4*, identified to promote cancer development, are upregulated in transition cells comparing to stable cells at 6W PTI. This also suggested the cellular behaviors of transition cells are different from that of stable cells even at the same sample collecting time point.

**Fig. 3.**
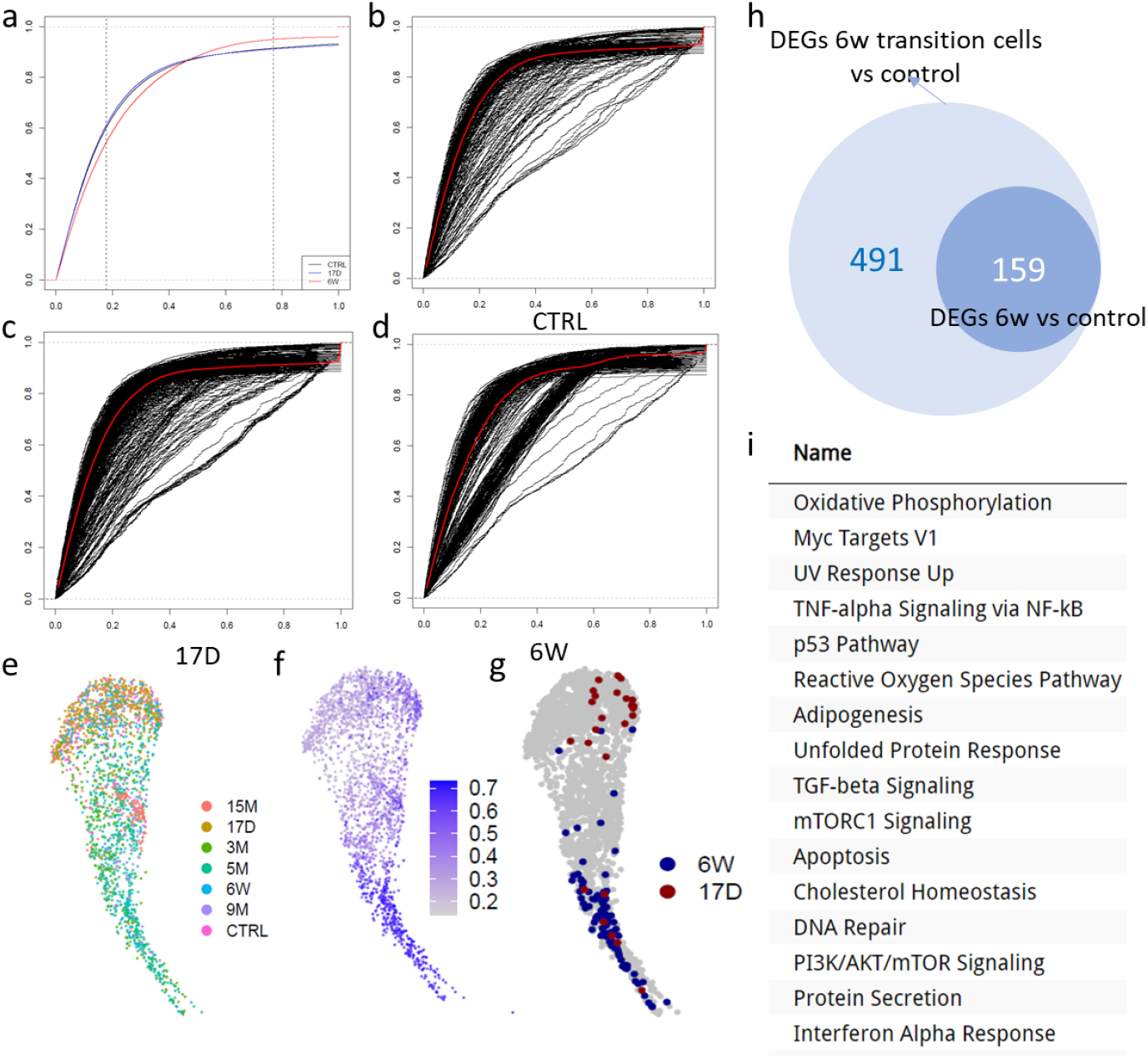
The distribution of gene pair-wise Pearson’s correlation coefficients illustrates ADM (Acinar ductal metaplasia) process through transition cells. **a-d**, The distribution of gene Pearson’s correlation are different across early stages of ADM process. The eCDF is different both between groups overall (**a**, black: control; blue: 17D PTI; red: 6W PTI) and within groups (**b**, control, **c**, 17D and **d**, 6W PTI). **e-f**, UMAP plot of acinar cells colored by **e**, transition index **f**, sample collecting time. **g**, Transition cells at 17D and 6W PTI highlighted on the UMAP. **h**, Number of DEGs found by transition cells and all cells 6W PTI comparing to the control group. **i**, HALLMARK pathway enrichment using 491 DEGs found only by transition cells.

To further investigate how transition cells can help us understand the early events of PDAC, we select 17 transition cells whose gene expressions are clustered closer to most of the acinar cells in the control group (**Fig. 4a**). Though there was no DEGs that could be found for these 17 transition cells when comparing the mean expressions with the control group, the gene pair-wise Pearson’s correlations of 5 genes are significantly increased in the transition cells (Wilcoxon test; p-value < 0.01). The expression level of these genes was quite low at early stage but were upregulated at later time points (**Fig. 4c**), suggesting the potential capability of using genes found by transition cells as early diagnosis signals.

**Fig. 4.**
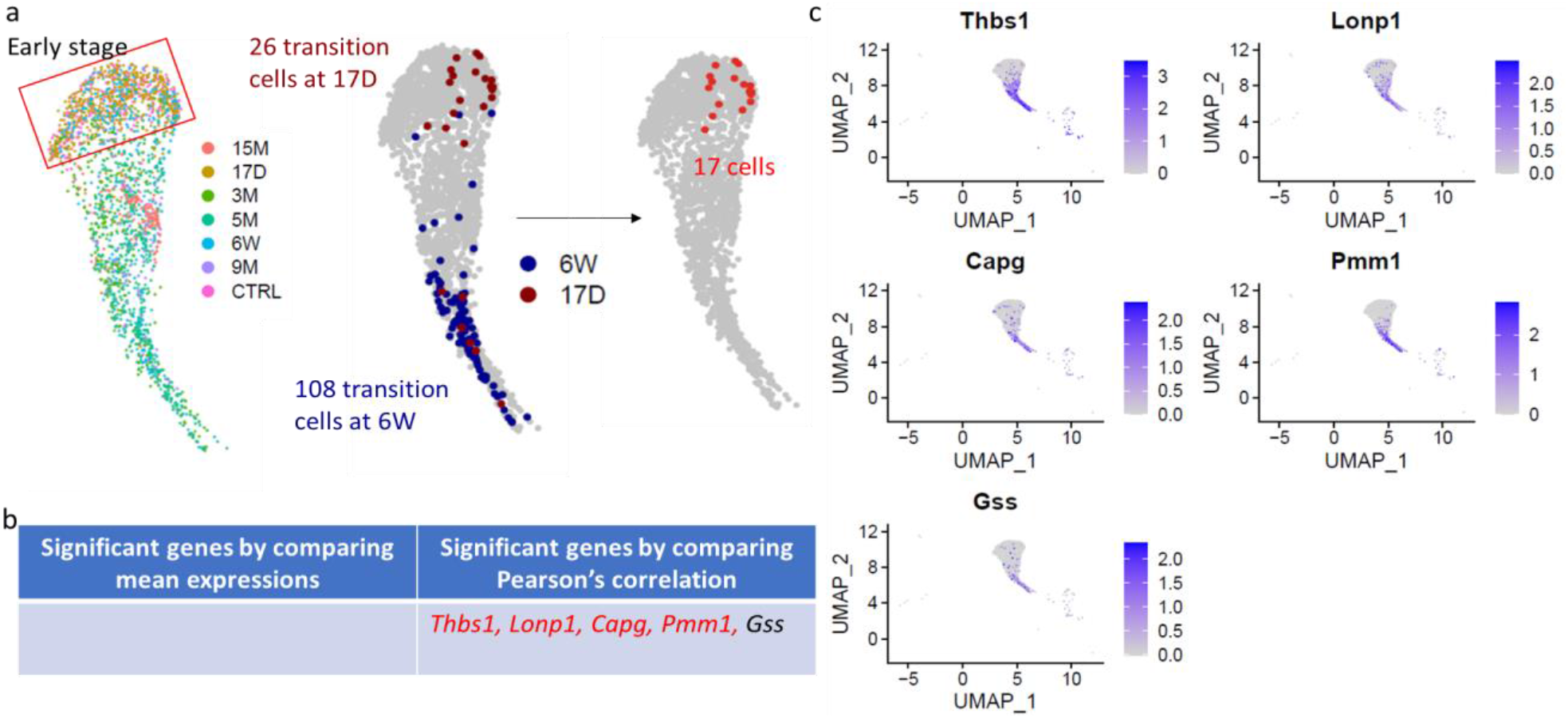
Transition cells and their corresponding significant genes at early stage of ADM phase can indicate the expression level at later time points. **a**, Transition cells at early stage of ADM phase. **b**, Significantly expressed genes found by Pearson’s correlation and mean expression when comparing transition cells at 17D PTI and control. (Red: differentially expressed at 6W PTI) **c**, The expression level of significant genes found by Pearson’s correlation.

## Discussions

Single-cell RNA-seq enables a high-resolution measurement of the dynamics during developmental processes. However, current analytical tools are deficient to study these dynamics and state transitions. Here, we propose a metric based on gene pair-wise Pearson’s correlation coefficients to quantify the transition cells and better understand the developmental processes. Transition cells are heterogenous and can imply distinct cell fates. Transition state pancreatic metaplastic cells at 3M PTI indicate different evolving directions of metaplastic cells. Moreover, transition cells identified by Pearson’s correlations reflect the alteration of the gene regulations underlying comparing to cells in the stable states, thus can give us opportunities to investigate the subtle changes during developmental processes. Taken together, our study bridged together dynamics systems theory with single cell RNA-seq, proposed a simple metrics and the analytical framework that can advance understanding molecular dynamics during both normal and abnormal developmental processes, and can potentially be applied to diagnosis, prognosis and therapeutics of diseases.

## Methods

### Single-cell RNA-seq datasets

Both mouse muscle cell regeneration and PDAC mouse model single-cell RNA-seq datasets were obtained from previous publications. The PDAC dataset were downloaded through GEO Series accession number GSE141017. And the mouse muscle cell regeneration dataset were downloaded at https://datadryad.org/stash/dataset/doi:10.5061%2Fdryad.t4b8gtj34

### Cell type annotation

Cell types were determined based on the original publications. Briefly, PDAC dataset has the public available metadata containing cluster numbers with cell barcodes. In the original publication, they provided a relation between cell types and the cluster number. We annotated cells by mapping the cluster number of each cell with the cell types according to the publication. Mouse muscle cell regeneration dataset makes the cell type information available in public repositories. We annotate committed progenitors and myoblasts as transition cells and others as stable cells according to the original publication and canonical markers.

### Analysis framework

The analysis framework was shown in **Fig. 5**. Briefly, single-cell RNA-seq data was normalized and scaled using Seurat (v4.0.0). Meta cells were generated by combining nearest neighbors in PCA dimensions to eliminate dropouts. Oscillating genes were removed using Oscope (v1.26.0) and top 100 most variable genes were selected for calculating gene pair-wise Pearson’s correlation based on nearest 200 neighbors in the PCA dimensions.

**Fig. 5.**
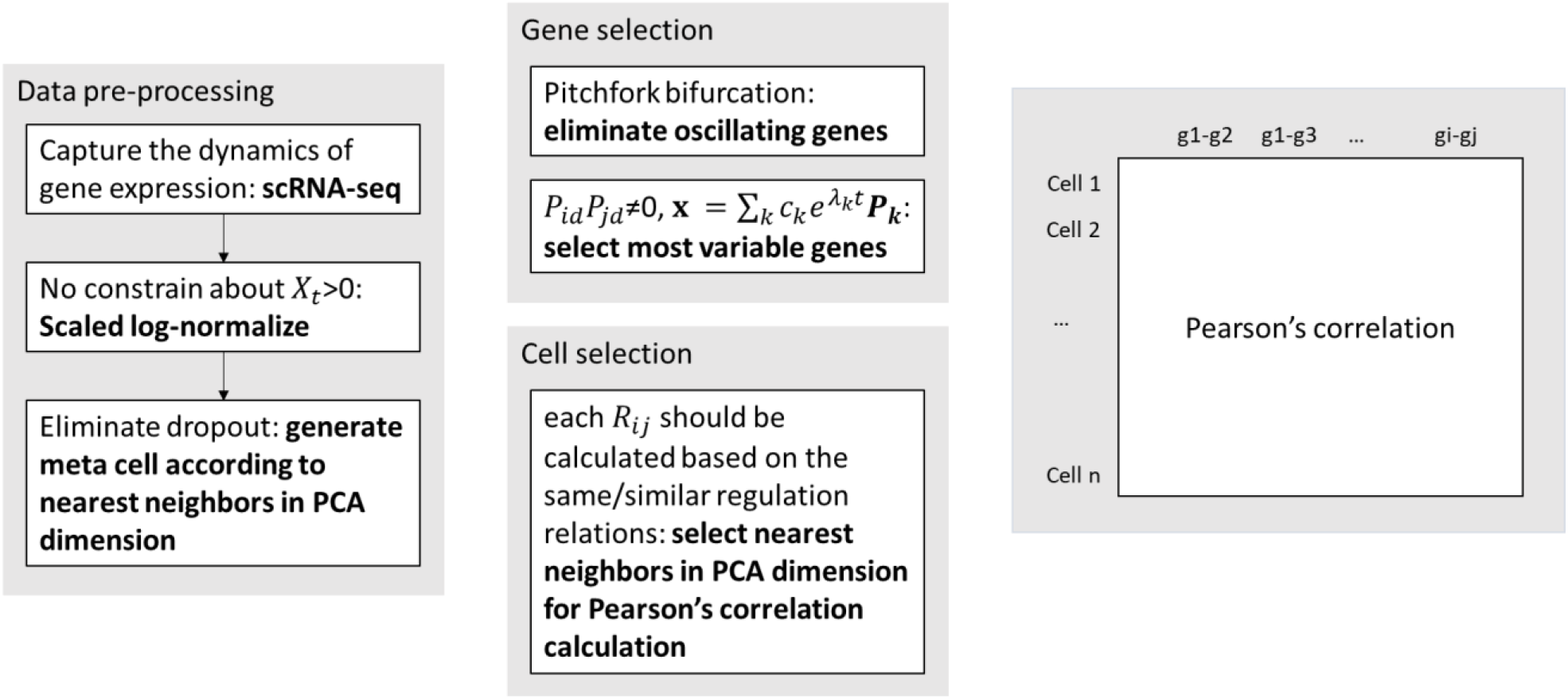
Analysis framework. Meta cells are generated through normalized and scaled scRNA-seq by combining their nearest neighbors to eliminate dropout issues. Most variable non-oscillating genes are selecting for computing gene pair-wise Pearson’s correlations.

### Transition index

Transition index was defined to quantify the possibility that a cell to be a transition cell according to the distribution of gene pair-wise Pearson’s correlations of the cell. We first found the maximal difference of eCDF of Pearson’s correlations between the reference group and group of interest, and then used the percentage of gene pairs within this range as the transition index.

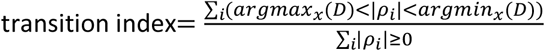

## Reference

Brackston, R. D., Lakatos, E., & Stumpf, M. P. H. (2018). Transition state characteristics during cell differentiation. PLoS Computational Biology, 14(9). https://doi.org/10.1371/JOURNAL.PCBI.1006405

Cardoso-Moreira, M., Sarropoulos, I., Velten, B., Mort, M., Cooper, D. N., Huber, W., & Kaessmann, H. (2020). Developmental Gene Expression Differences between Humans and Mammalian Models. Cell Reports, 33(4), 108308. https://doi.org/10.1016/J.CELREP.2020.108308

Daun, S., Rubin, J., Vodovotz, Y., & Clermont, G. (2008). EQUATION-BASED MODELS OF DYNAMIC BIOLOGICAL SYSTEMS. Journal of Critical Care, 23(4), 585. https://doi.org/10.1016/J.JCRC.2008.02.003

Freedman, S. L., Xu, B., Goyal, S., & Mani, M. (2022). A dynamical systems treatment of transcriptomic trajectories in hematopoiesis. BioRxiv, 2021.05.03.442465. https://doi.org/10.1101/2021.05.03.442465

Hwang, B., Lee, J. H., & Bang, D. (2018). Single-cell RNA sequencing technologies and bioinformatics pipelines. Experimental & Molecular Medicine 2018 50:8, 50(8), 1–14. https://doi.org/10.1038/s12276-018-0071-8

Ioannis Stefanou, & Jean Sulem. (2021). Instabilities Modeling in Geomechanics. In John Wiley & Sons. https://books.google.com/books?hl=en&lr=&id=q_ArEAAAQBAJ&oi=fnd&pg=PA31&dq=bifurcation+theory+ode&ots=Cj5baFqsmB&sig=euCeZG3KtPbQ4czzXqt4ZSmvUuQ#v=onepage&q=bifurcation%20theory%20ode&f=false

Kreutz, C. (2020). A New Approximation Approach for Transient Differential Equation Models. Frontiers in Physics, 0, 70. https://doi.org/10.3389/FPHY.2020.00070

Lange, M., Bergen, V., Klein, M., Setty, M., Reuter, B., Bakhti, M., Lickert, H., Ansari, M., Schniering, J., Schiller, H. B., Pe’er, D., & Theis, F. J. (2022). CellRank for directed single-cell fate mapping. Nature Methods 2022 19:2, 19(2), 159–170. https://doi.org/10.1038/s41592-021-01346-6

Ma, Z., Lytle, N. K., Chen, B., Jyotsana, N., Novak, S. W., Cho, C. J., Caplan, L., Ben-Levy, O., Neininger, A. C., Burnette, D. T., Trinh, V. Q., Tan, M. C. B., Patterson, E. A., Arrojo e Drigo, R., Giraddi, R. R., Ramos, C., Means, A. L., Matsumoto, I., Manor, U., … DelGiorno, K. E. (2022). Single-Cell Transcriptomics Reveals a Conserved Metaplasia Program in Pancreatic Injury. Gastroenterology, 162(2), 604-620.e20. <https://doi.org/10.1053/J.GASTRO.2021.10.027/ATTACHMENT/2564CDDB-D03F-4EAD-8CF2-C61526A942AB/MMC8.MP4 >

McKellar, D. W., Walter, L. D., Song, L. T., Mantri, M., Wang, M. F. Z., de Vlaminck, I., & Cosgrove, B. D. (2021). Large-scale integration of single-cell transcriptomic data captures transitional progenitor states in mouse skeletal muscle regeneration. Communications Biology 2021 4:1, 4(1), 1–12. https://doi.org/10.1038/s42003-021-02810-x

Mulas, C., Chaigne, A., Smith, A., & Chalut, K. J. (2021). Cell state transitions: definitions and challenges. Development (Cambridge), 148(20). https://doi.org/10.1242/DEV.199950/272516 Pancreatic Cancer Prognosis | Johns Hopkins Medicine. (n.d.). Retrieved May 11, 2022, from https://www.hopkinsmedicine.org/health/conditions-and-diseases/pancreatic-cancer/pancreatic-cancer-prognosis

Reyes-Castellanos, G., Masoud, R., & Carrier, A. (2020). Mitochondrial Metabolism in PDAC: From Better Knowledge to New Targeting Strategies. Biomedicines 2020, Vol. 8, Page 270, 8(8), 270. https://doi.org/10.3390/BIOMEDICINES8080270

Rodriguez-Esteban, R., & Jiang, X. (2017). Differential gene expression in disease: A comparison between high-throughput studies and the literature. BMC Medical Genomics, 10(1), 1–10. <https://doi.org/10.1186/S12920-017-0293-Y/FIGURES/6 >

Schlesinger, Y., Yosefov-Levi, O., Kolodkin-Gal, D., Granit, R. Z., Peters, L., Kalifa, R., Xia, L., Nasereddin, A., Shiff, I., Amran, O., Nevo, Y., Elgavish, S., Atlan, K., Zamir, G., & Parnas, O. (2020). Single-cell transcriptomes of pancreatic preinvasive lesions and cancer reveal acinar metaplastic cells’ heterogeneity. Nature Communications 2020 11:1, 11(1), 1–18. https://doi.org/10.1038/s41467-020-18207-z

Wang, Z., Feng, X., & Li, S. C. (2019). SCDevDB: A database for insights into single-cell gene expression profiles during human developmental processes. Frontiers in Genetics, 10(SEP), 903. https://doi.org/10.3389/FGENE.2019.00903/BIBTEX

Zhou, P., Wang, S., Li, T., & Nie, Q. (2021). Dissecting transition cells from single-cell transcriptome data through multiscale stochastic dynamics. Nature Communications 2021 12:1, 12(1), 1–15. https://doi.org/10.1038/s41467-021-25548-w

